# Structural classification of proteins based on the computationally efficient recurrence quantification analysis and horizontal visibility graphs

**DOI:** 10.1101/2020.10.23.350736

**Authors:** Michaela Areti Zervou, Effrosyni Doutsi, Pavlos Pavlidis, Panagiotis Tsakalides

## Abstract

**Motivation:** Protein structure prediction is one of the most significant problems in bioinformatics, as it has a prominent role in understanding the function and evolution of proteins. Designing a computationally efficient but at the same time accurate prediction method remains a pressing issue, especially for sequences that we cannot obtain a sufficient amount of homologous information from existing protein sequence databases. Several studies demonstrate the potential of utilizing chaos game representation (CGR) along with time series analysis tools such as recurrence quantification analysis (RQA), complex networks, horizontal visibility graphs (HVG) and others. However, the majority of existing works involve a large amount of features and they require an exhaustive, time consuming search of the optimal parameters. To address the aforementioned problems, this work adopts the generalized multidimensional recurrence quantification analysis (GmdRQA) as an efficient tool that enables to process concurrently a multidimensional time series and reduce the number of features. In addition, two data-driven algorithms, namely average mutual information (AMI) and false nearest neighbors (FNN), are utilized to define in a fast yet precise manner the optimal GmdRQA parameters.

**Results:** The classification accuracy is improved by the combination of GmdRQA with the HVG. Experimental evaluation on a real benchmark dataset demonstrates that our methods achieve similar performance with the state-of-the-art but with a smaller computational cost.

**Availability:** The code to reproduce all the results is available at https://github.com/aretiz/protein_structure_classification/tree/main.

**Supplementary information:** Supplementary data are available at *Bioinformatics* online.

## 1 Introduction

Protein structure prediction is one of the most important and challenging issues in computational biology. Obtaining knowledge of protein function and regulation is highly important and useful in medicine and biotechnology (Noble *et al.*, 2004), especially for drug design, enzymes composition and interpretation of disease related phenotypes. Proteins function properly when they adopt their final and stable shape which is also known as tertiary structure or fold. According to the Anfinsen’s dogma the protein folding is mainly determined by its amino acid sequence (Anfinsen, 1973). Based on their folding patterns, proteins can be classified into four structural categories, namely, (i) all-*α*, where the structural domains are mainly composed of *α*-helices and a small amount of *β*-strands (a.k.a. *β*-sheets), (ii) all-*β*, that is mostly formed by *β*-strands and a few isolated *α*-helices, (iii) *α* + *β*, forming *α*-helices and mostly anti-parallel *β*-strands, and (iv) *α/β* consisting of *α*-helices and almost all parallel *β*-strands (Levitt and Chothia, 1976). The last few decades, the accelerated evolution of genomics has lead to a substantial volume of amino acid sequence data of proteins. Therefore, an emerging need of efficient architectures for protein structure classification has arisen. Along these lines, a plethora of machine learning based algorithms has been developed for protein structural class prediction. Some extensively utilized methods for the representation of protein samples via feature extraction are the amino and pseudo-amino acid composition (Nakashima *et al.*, 1986; Chou, 2001), the identification of binding motifs in protein–protein interactions (Guharoy and Chakrabarti, 2007), the PSI-BLAST profile (Liu *et al.*, 2010), and the predicted secondary structure information (Liu and Jia, 2010), among others. However, the performance of these techniques suffers when low-similarity proteins are encountered. Thus, significant effort has been put to improve the prediction accuracy for proteins for which we cannot obtain a sufficient amount of homologous information (Yang *et al.*, 2008; Zhang *et al.*, 2012; Liang *et al.*, 2015; Wang *et al.*, 2015; Yu *et al.*, 2017; Zhu *et al.*, 2019; Apurva and Mazumdar, 2020).

Numerous studies (Yang *et al.*, 2009, 2010; Olyaee *et al.*, 2016; Jiang *et al.*, 2019; Olyaee *et al.*, 2016) demonstrate the potential of transforming the amino acid sequence into a time series and then, utilizing powerful time series analysis techniques such as Recurrence Quantification Analysis (RQA) (Eckmann *et al.*, 1987), Horizontal Visibility Graphs (HVG) (Lacasa *et al.*, 2008) or a combination of both, for extracting meaningful information from the data. The general procedure for protein structure classification is briefly depicted in Fig.1. According to the scheme, every amino acid in the protein sequence is initially predicted as one of three secondary structural elements, namely H (helix), E (strand) and C (coil) using the PSI-PRED (Jones, 1999) tool and then, by employing the Chaos Game Representation (CGR) (Jeffrey, 1990) technique it is possible to generate the time-series which are processed to create the set of features. Although the aforementioned studies lead to high-precision results, their enhanced performance comes at the expense of time and memory complexity as well as with an inefficient processing of the multidimensional data. This is mainly due to the fact that (i) a pair of coordinates has to be processed separately, resulting in a large number of features and (ii) the fine-tuning of the RQA parameters for each time series is highly time demanding.

**Fig. 1:**
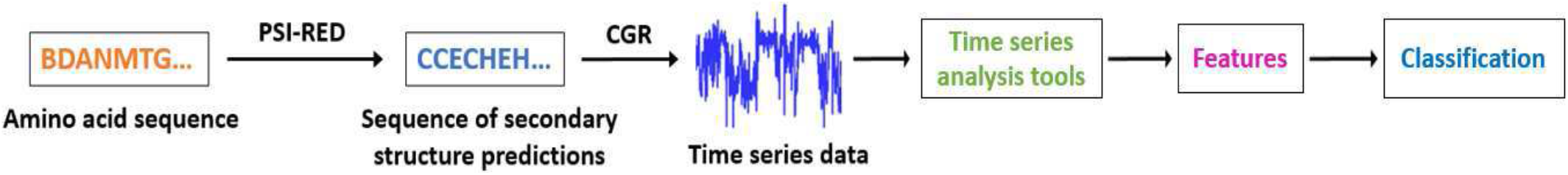
General protein structure prediction scheme.

To overcome these limitations, this work processes directly on the two-dimensional time series by introducing (i) the utilization of the Generalized multidimensional Recurrence Quantification Analysis (GmdRQA) (Zervou *et al.*, 2019) as a sophisticated, non-linear analysis tool of multidimensional time series data, and (ii) a data-driven estimation of the RQA parameters employing the Average Mutual Information (AMI) (Fraser and Swinney, 1986) and the False Nearest Neighbors (FNN) (Kennel *et al.*, 1992) methods. The aforementioned mechanisms enable to exploit both the intra- and inter-data correlations among the two time series in an automated fashion, resulting in lower time and memory complexity. The contributions of this paper are summarized below:

i. The utilization of GmdRQA greatly reduces the number of features as it is applied directly to the two-dimensional time series.
ii. The introduction of a data-driven parameter selection employing the FNN and AMI algorithms.
iii. The design of a novel feature extraction scheme consisting of HVG and the Data-Driven unidimensional RQA (DD-RQA) or the Data-Driven GmdRQA (DD-GmdRQA) frameworks that enable the discovery of representative patterns, increasing the overall accuracy of the protein structure prediction.
iv. The significant reduction of the computational complexity of both the feature selection and the classification processes.

The rest of the paper is organized as follows: Section 2 is a background on the most notable methods that have been used in the literature for the generation of time series from an amino acid sequence and gives details on the proposed GmdRQA architectures that consist of two data-driven parameter-tuning algorithms, namely AMI and FNN. Section 3 introduces the proposed time delay embedding parameter selection architectures, the classification procedure and the performance of the proposed architectures in terms of classification accuracy, time complexity and feature multitude. Finally, section 4 draws the conclusion of this work and gives directions for future extensions.

## 2 Methods

The purpose of this section is to briefly introduce the most significant mechanisms that have been used in the literature in order to (i) transform the amino acid sequence into time series and (ii) extract informative features for an accurate protein structure prediction using machine learning algorithms. Specifically, the PSI-PRED and the CGR methods are first introduced as they are employed to convert an amino-acid sequence into a time series. Then, two time series analysis techniques, the RQA and the HVG, are described in details to extract information-rich characteristics.

### 2.1 PSI-blast based secondary structure PREDiction

Each protein is a long chain called polypeptide or polymer, which is formed when several monomers, known as amino acids, are joined together. Only 20 amino acids are known as proteinogenic, meaning they participate in the synthesis of a protein primary structure. In the literature, there are several works (Yang *et al.*, 2010; Olyaee *et al.*, 2016; Jiang *et al.*, 2019) that instead of dealing with the protein primary structure they use the PSI-blast based secondary structure PREDiction (PSIPRED) tool that predicts the role of each amino acid in the protein secondary structure. Particularly, PSI-PRED transforms the initial amino acid sequence to a sequence of equal length that now consists of only three states that describe its secondary structure, namely coils (C), strands (E) and helices (H). This simplification reduces the dimensionality of our data from 20 amino acids to three structural elements, thus easing the overall computational complexity. Hence, in this work, we have decided to use as input data the prediction of the protein secondary structure.

### 2.2 Time Series Generation via Chaos Game Representation

In order to transform the unidimenisional sequence of characters into a two-dimensional time series, chaos game representation (CGR) is employed. CGR was first proposed as a scale-independent representation of genomics (Jeffrey, 1990; Almeida *et al.*, 2001). In essence, CGR is able to graphically represent the sequence while preserving its original structure. Specifically, a sequence is represented in a unit equilateral triangle. Its three vertices refer to the three secondary structure types namely helix (H), coils (C), and strands (E), with *xy*-plane coordinates (0,0),(0.5,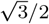) and (1,0), respectively. The CGR graph, as shown in Fig. 2 (Left) is obtained through the following procedure: Initially, the triangle centroid 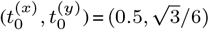 is defined and then, the 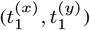 coordinates of the first element of the sequence are calculated as the halfway distance point between the centre of the triangle and the vertex representing this element. Accordingly, the remaining consecutive elements in the secondary structure sequence are plotted as the midpoint between the previous plotted point and the vertex representing the element being plotted as follows,

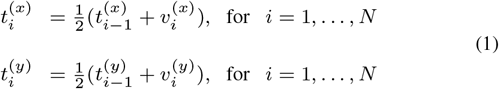

where 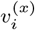 and 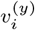 are respectively the *x* and *y* coordinates of the vertex corresponding to the *i*^th^ secondary structure element of a protein consisting of *N* amino-acids. Finally, as depicted in Fig. 2 (Middle) and Fig. 2 (Right), the CGR graph is decomposed into two time series that consist accordingly of the *x* and *y* coordinates so that 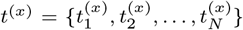 and 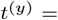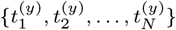.

**Fig. 2:**
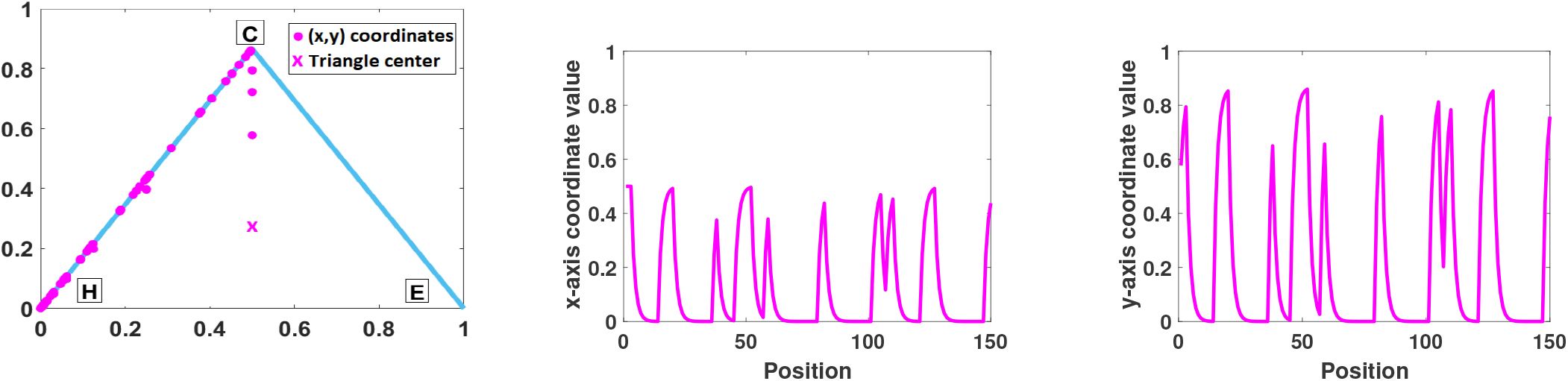
Left: Chaos Game Representation of protein’s 1ASH predicted secondary structure (’*CCCHH… HHCCC*’) based on the three secondary structural elements, C (coil), E (strand) and H (helix). This graph indicates that the sequence consist of only coil and helix elements. Middle: This time series represent the x-coordinates of the points. Right: This time series represent the x-coordinates of the points.

### 2.3 Time Series Analysis

#### 2.3.1 Recurrence Quantification Analysis

The recurrence quantification analysis (RQA) (Eckmann *et al.*, 1987) is exploited to perform a sophisticated non-linear analysis of the time series data. RQA is capable of treating non-stationary and short data series, as it comprises a set of appropriate quantitative measures for the analysis of recurrences, typically small-scale structures. As a result, RQA enables the detection of critical transitions in the system’s dynamics (e.g. deterministic, stochastic, random). More specifically, a recurrence plot (RP) is derived depicting those times at which a state of a dynamical system recurs. In particular, the recurrence of a state that occurs at time *i* and at a different time *j* is represented within a two-dimensional square matrix with ones (recurrence) and zeros (non-recurrence), where both axes are time axes. In other words, RPs reveal all the times when the phase space trajectory of the dynamical system visits roughly the same area in the phase space. To this end, RPs enable the investigation of an *m*-dimensional phase space trajectory through a two-dimensional representation of its recurrences.

Given a time series of length *N*, 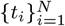, a phase space trajectory can be reconstructed via time-delay embedding,

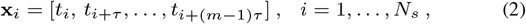

where *m* is the embedding dimension, *τ* is the time delay, and *N*_*s*_ = *N* −(*m*−1)*τ* is the number of states. Having constructed a phase space representation, an RP is defined as follows,

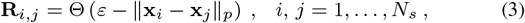

where x_*i*_, x_*j*_ ∈ ℝ^*m*^ are the states, *ε* is a threshold, ∥ · ∥_*p*_ denotes a general *ℓ*_*p*_ norm, and Θ(·) is the Heaviside step function, whose discrete form is defined by

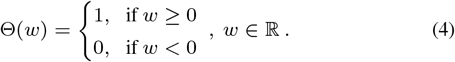

The resulting matrix **R** exhibits the main diagonal, **R**_*i,i*_ = 1, *i* = 1*,…, N*_*s*_, also known as the *line of identity* (LOI). Typically, several linear (and/or curvilinear) structures appear in RPs, which give hints about the time evolution of the high-dimensional phase space trajectories. A major advantage of RPs is that they can also be applied to rather short and even non-stationary data. The visual interpretation of RPs, which is often difficult and subjective, is enhanced by means of several numerical measures for the quantification of the structure and complexity of RPs (Zbilut and Webber, 1992). These quantification measures provide a global picture of the underlying dynamical behavior during the entire period covered by the data. This work utilizes 10 of the RQA quantitative measures (Marwan *et al.*, 2007 that can be found in the Supplementary data file.

#### 2.3.2 Horizontal Visibility Graph

In recent years, complex network theory has been popularized in the analysis of biological problems (Zhao *et al.*, 2018). A simple and fast computational method, known as horizontal visibility graph (HVG) (Lacasa *et al.*, 2008), maps time series into graphs. HVG is invariant under affine transformations of the series data and its main focus lies on the time series structural properties (periodicity, fractality, etc.). Specifically, let a time series 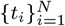. The HVG algorithm assigns each sample point *t*_*i*_ as a node *n*_*i*_ of a graph *G*. Then, two nodes *n*_*i*_ and *n*_*j*_ in the network are connected if the geometrical rule in (5) is satisfied,

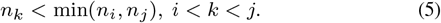

In essence, two nodes *n*_*i*_ and *n*_*j*_ share an edge when a horizontal line can be drawn among them without intersecting, in terms of magnitude, any intermediate node. In this study, the resulted horizontal visibility graph *G* = (*V, E*), *N* = |*V*|, *M* = |*E*|, with *N* and *M* being the number nodes and edges respectively, is undirected, unweighted and connected. The graph properties are represented by the measures described in the Supplementary data file that are later employed for classification purposes.

### 2.4 Dataset Description

This work employs the 25PDB datasetthat includes 1673 proteins of varying length with 25% sequence homology. The length wise distribution of the protein sequences is comparable for the all-*α*, all-*β* and *α* + *β* folds, whereas for the *α/β* fold the length is observed to be generally higher. Particularly, in the all-*α*, all-*β* and *α* + *β* folds, there is a higher proportion of small protein sequences with less than 100 residues compared to the *α/β* fold. On the other hand, the number of sequences that consist of more than 300 residues is higher for the *α/β* and *α* + *β* folds against all-*α* and all-*β* folds. The proteins are categorised based on their structural class as following: 443 proteins belong to the all-*α*, 443 to the all-*β*, 346 to the *α/β* and 441 to *α* + *β* fold.

### 2.5 Generalized Multidimensional Recurrence Quantification Analysis

Multidimensional recurrence quantification analysis Wallot *et al.*, 2016 extracts the underlying dynamics of the system by mapping the time series in a higher dimensional phase space of trajectories by constructing state vectors u_*i*_ via time delay embedding. The generalized multidimensional recurrence quantification analysis (GmdRQA) framework transforms state vectors u_*i*_ into state matrices *X*_*i*_ to represent the time-delay embedding (Zervou *et al.*, 2019). This is due to the fact that state matrices are considered more appropriate for describing multidimensional signals from a mathematical perspective, enabling them to model the correlations not only within a signal but also between different signals. More specifically, given a multidimensional time series 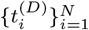, where *D* stands for the data dimensionality, the corresponding phase space representation is reconstructed as follows,

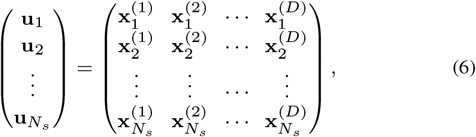

where 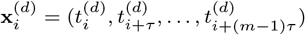, *i* = 1*,…, N*_*s*_, *d* = 1*,…, D*, *m* being the embedding dimension, *τ* the delay, and *N*_*s*_ = *N* − (*m* − 1)*τ* the number of states. The state vectors u_*i*_ can be transformed into state matrices of the form

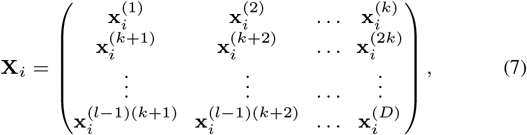

where 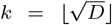, *l* = ⌊*D/k*⌋ and *i* = 1*,…, N*_*s*_. Subsequently, the Generalized multidimensional Recurrence Plot (GmdRP) is defined according to (3). Having constructed the corresponding recurrence plot of the multidimensional system, the features described in the Supplementary data file can also be employed.

### 2.6 Estimation of embedding parameters

Identifying the optimal RQA/GmdRQA parameters for the reconstruction of the phase space is extremely crucial. To the best of our knowledge, parameter tuning is usually done in a grid search manner which is time demanding. In the herein work, the estimation of the embedding parameters is performed in a Data-Driven (DD) fashion where the AMI and the FNN methods are utilized to evaluate the optimal time delay *τ* and minimal sufficient value of the embedding dimension *m*, respectively.

#### 2.6.1 Average Mutual Information Algorithm

The Average Mutual Information (AMI) (Fraser and Swinney, 1986) is a measure of nonlinear correlation between the given signal 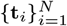 and a time delayed version of this signal by *τ* samples 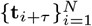 and is expressed as,

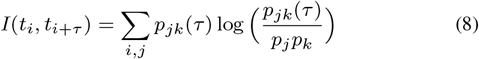

where *p*_*j*_ is the probability that *t*_*i*_ is in bin *j* of the histogram constructed from the data points in *t*, and *p*_*jk*_(*τ*) is the probability that *t*_*i*_ is in bin *j* and *t*_*i*+*τ*_ is in bin *k*.

Determining a proper value for *τ* implies that the coordinates of the phase space embedded signal will be maximally independent. As proposed by (Fraser and Swinney, 1986, this is guaranteed by choosing as the optimal value for the time lag *τ* the position of the first minimum of *I*(*t*_*i*_, *t*_*i*+*τ*_). Nonetheless, it is possible that the AMI function does not acquire a local minimum. Therefore, Kantz and Schreiber, 2004 introduced a criterion where the optimal value is considered to be the lowest value of *τ* for which the AMI function descents below the value 1/e, *e* ≈ 2.71. Furthermore, in this study, the max lag *τ* as well as the number of bins for calculating the histogram are set to 10.

#### 2.6.2 False Nearest Neighbor Algorithm

The embedding dimension *m* is an estimate of the dimensionality of the dynamics of the time series. The False Nearest Neighbor (FNN) (Kennel *et al.*, 1992) method is based on the assumption that two points that are close in the sufficient embedding dimension should continue to be close as the dimension increases. A criterion for recognizing embedding errors is a considerable increase in the distance between two neighbouring points while moving from dimension *m* to *m* + 1.

Specifically, given an embedded time series in a *m*-dimensional phase space with a time delay *τ* and two of its coordinate vectors x_*i*_ and x_*j*_ that are adjacent at a time instance, the squared Euclidean distance between them when moving from *m* into (*m* + 1) dimensions is,

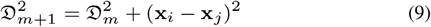

where 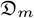 is the Euclidean distance between x_*i*_ and x_*j*_ and is defined as,

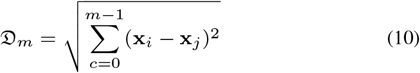

If the one-dimensional time series is already properly embedded in *m* dimensions, then the distance 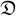 between x_*i*_ and its nearest neighbor x_*j*_ should not appreciably change by some distance criterion 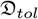 so that 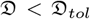. Moreover, the distance of the nearest neighbor when embedded into the next higher dimension should be less than some criterion *A*_*tol*_ such that 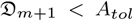. According to Wallot and Mønster, 2018 the next settings of the thresholds 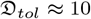 and *A*_*tol*_ = 2 are recommended. The procedure is repeated for the nearest neighbor of each coordinate vector until one of the stopping criteria is met. In particular, the optimal *m* is reached when i) FNN drops to 0, or ii) subsequent embeddings have the same number of false neighbors which implies that their difference is less than a threshold *T*_*tol*_ = 2, or iii) the point before which the number of FNNs starts to increase again.

## 3 Results

This section initially describes the experimental evaluation for choosing the optimal set of parameters, *m*, *τ* and *ε*, of the herein proposed RQA parameter selection schemes. Then the performance of each proposed architecture is compared to state-of-the-art in terms of overall classification accuracy, feature multitude, and running time complexity. It is important to note that in this work the classification procedure is not considered for the measurement of computational complexity. All the experiments are implemented in MATLAB, on a desktop computer equipped with a CPU processor (Intel Core i5-4590) clocked at 3.30GHz, and a 8 GB RAM.

### 3.1 RQA Parameter Selection

The goal of this section is to show that the GmdRQA is capable of *reducing* the number of features by half and *increasing* the computational efficiency of the system while achieving the same prediction accuracy. In particular, a parameter selection based on Grid Search (GS) is evaluated as it is the only technique suggested in the literature for the protein prediction problem. In addition to GS, two more case studies are proposed and evaluated: (i) the embedding dimension *m* and time delay *τ* parameters are found for each protein in a Data-Driven (DD) fashion when AMI and FNN are employed, and (ii) the Most Frequent (MF) *m* and *τ* values are computed as the product of a statistical analysis of the optimal set of parameters. These three parameter selection approaches are employed by RQA and GmdRQA, resulting in 6 different algorithms. Last but not least, HVG is also combined with each one of the 6 aforementioned algorithms. The general framework of the proposed protein structure prediction architecture is depicted in Fig. 3.

**Fig. 3:**
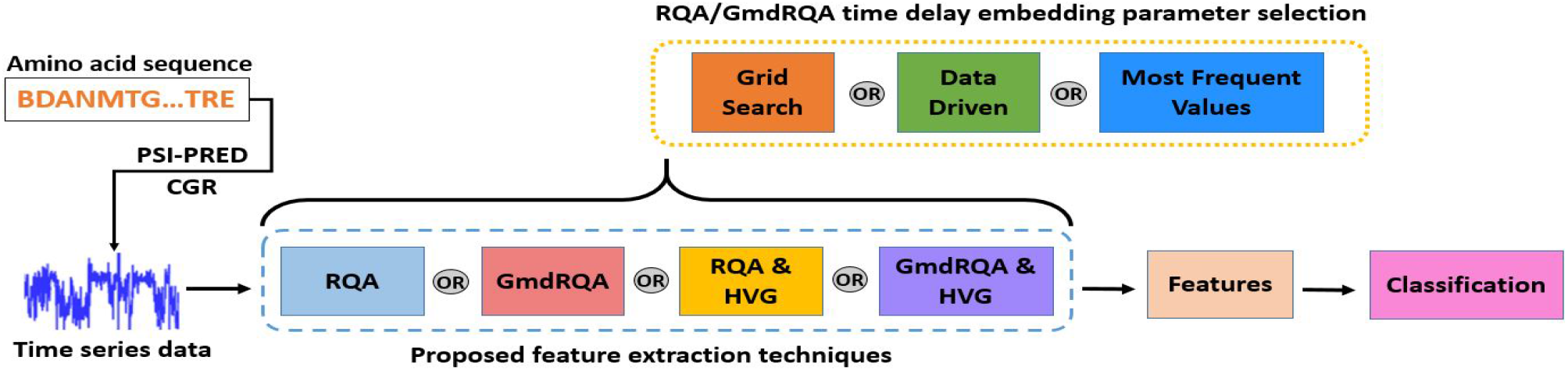
General proposed RQA time delay embedding parameter selection scheme.

#### 3.1.1 Grid Search Parameter Tuning

The performance of GS-RQA and GS-GmdRQA schemes is initially evaluated when the range of embedding dimension *m* and time delay *τ* varies between 1 and 8. In the case that both *m* and *τ* are equal to 8, the phase space cannot be constructed for small proteins, hence no results are reported and the grid search procedure is terminated. To assess the classification accuracy of GS-RQA and GS-GmdRGA, we generated 30 randomly shuffled datasets and for each dataset the classification accuracy is reported for all combinations of *m* and *τ* for every individual shuffling. The pair that is reported to most frequently achieve the highest accuracy among all the different random data shuffles is employed as the optimal. Based on this procedure, the optimal parameters for GS-RQA are *m* = 4 and *τ* = 1 when FDA is employed and *m* = 2 and *τ* = 4 for SVM. On the other hand, for GS-GmdRQA, the optimal parameters for FDA are *m* = 2 and *τ* = 4 while for FDA the optimal parameters are *m* = 5 and *τ* = 1. The implementation of GS-RQA and GS-GmdRQA for the rest of this work uses the herein evaluated optimal parameters.

#### 3.1.2 Data-driven Parameter Tuning

The following case study involves the data-driven fine-tuning time delay embedding parameter selection. In this scenario, the optimal set of parameters is evaluated in an automated fashion. Particularly, the time delay embedding parameters are evaluated per protein using the FNN and AMI algorithms. However, in this work, for proteins of length lower than 45 residues, time delay embedding parameters are set to be *τ* = 1 and *m* = 2 that are the minimum values for both parameters. For the DD-RQA scheme, 20 features (10 features×2 dimensions) are extracted in total, whereas for the DD-GmdRQA architecture, the number of features is reduced to 10 since the two-dimensional time series is processed concurrently.

#### 3.1.3 Statistical Parameter Tuning

In the case of MF-RQA and MF-GmdRQA, the *m* and *τ* parameter values are selected as the most frequent values among the optimal sets of *m* and *τ* that derive when FNN and AMI are applied on each single protein included in the training phase to later avoid overfitting. Again, for proteins of length lower than 45 residues, the time delay embedding parameters are set as *τ* = 1 and *m* = 2 that are the minimum values for both parameters. The main difference between GS-RQA/GS-GmdRQA and MF-RQA/MF-GmdRQA is that for the grid search process (GS-RQA/GS-GmdRQA), *m* and *τ* assume the same value for both time series dimensions, whereas for the most frequent parameter searching (MF-RQA/MF-GmdRQA), *m* and *τ* are evaluated separately per dimension. Along these lines, concerning MF-RQA the most frequent values are (*m, τ*) = (4, 5) and (*m, τ*) = (2, 5) for the *t*_*x*_ and *t*_*y*_ time series, respectively. On the other hand, in MF-GmdRQA, (*m, τ*) = (2, 6) for both dimensions. For the rest of this work, the implementation of MF-RQA and MF-GmdRQA utilizes the aforementioned optimal parameters accordingly.

#### 3.1.4 Neighborhood ε

Following the empirical rule proposed by Krämer *et al.*, 2018 the neighborhood threshold *ε* can be chosen as a percentage of the distribution of all the pairwise distances that describe the phase space trajectory. Fig. 4 provides the average classification accuracy as a function of the neighborhood threshold *ε* for both the RQA and GmdRQA schemes using the FDA and SVM classifiers. As depicted, the average classification accuracy of GS-RQA and GS-GmdRQA plateaus for *ε* values above 30% for both classifiers for all study cases. As the focus of this study is to evaluate the influence of time and delay embedding parameters on the proposed architectures we select the same neighborhood percentage in all three models. Therefore, since a higher neighborhood percentage increases the computational cost, *ε* = 30% is selected as the minimum percentage that provides high classification results. The DD-RQA architecture performs its best when *ε* lies between 30%-50%. Thus, *ε* = 30% is chosen. In addition, for the case of DD-GmdRQA, it is clear that *ε* = 30% provides the highest accuracy over all percentages for both SVM and FDA. Finally, the performance of MF-RQA and MF-GmdRQA for FDA and SVM is better when *ε* lies between 20%-50% and 30%-50%, respectively. Consequently, we select *ε* = 20% for FDA and *ε* = 30% for SVM.

**Fig. 4:**
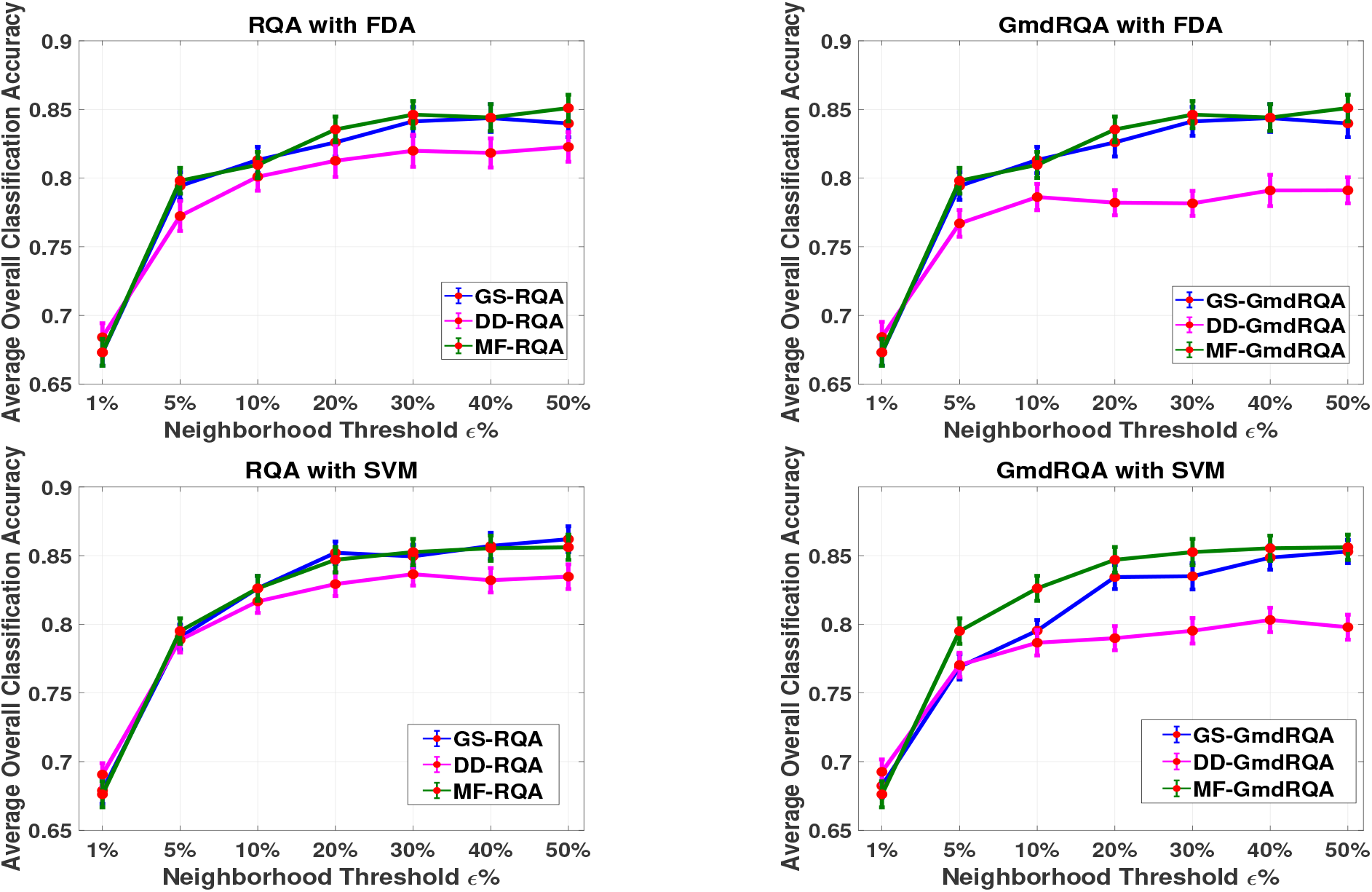
Mean classification accuracy as function of the percentage of the neighborhood threshold *ε* for RQA and GmdRQA with FDA (Top Left, Top Right) and SVM (Lower Left, Lower Right).

### 3.2 Classification

The first step in the classification process is to perform a z-score normalization of the feature matrix. Previous studies (Yang *et al.*, 2009, 2010; Olyaee *et al.*, 2016; Jiang *et al.*, 2019) recommend the leave-one-out cross-validation process. However, splitting the data into 70%-30% for training and testing respectively, yields nearly the same overall accuracy in considerably less time. Therefore, to reduce the running time complexity of the classification process, in this work the data are randomly split into 70%-30% for training and testing and the procedure is repeated 150 times. Along these lines, a Gaussian-kernel Support Vector Machine (SVM) as well as Fisher’s Linear Discriminant Analysis (FDA) algorithm are applied separately on the normalized feature matrix for discriminating between the four structural classes all-*α*, all-*β*, *α/β* and *α* + *β*. Concerning the SVM classifier, the regularization parameter *C* and kernel width parameter *γ* can take all positive values log-scaled in the range [10^−3^, 10^3^].Since each experiment is repeated 150 times, the average score of all the metrics mentioned above along with the respected standard deviation is reported.

Initially, the RQA and GmdRQA architectures are examined for each of the data-driven and the statistical time delay embedding parameter selection schemes (see sections 3.1.1 and 3.1.2). Particularly, Tables 1 and 2 indicate the performance of the proposed DD-RQA, DD-GmdRQA, MF-RQA and MF-GmdRQA feature extraction frameworks in terms of sensitivity, specificity and Overall Accuracy (OA) for SVM and FDA, respectively. As shown, proteins that belong to the *α*-fold and *β*-fold class are better predicted in most test cases for both SVM and FDA classifiers. Moreover, it is given that MF-RQA outperforms MF-GmdRQA in addition to DD-RQA and DD-GmdRQA. This leads to the conclusion that a more generalized approximation of the time-delay embedding parameters enhances the system’s ability to learn information-rich patterns that best capture the underlying data dynamics.

**Table 1.**
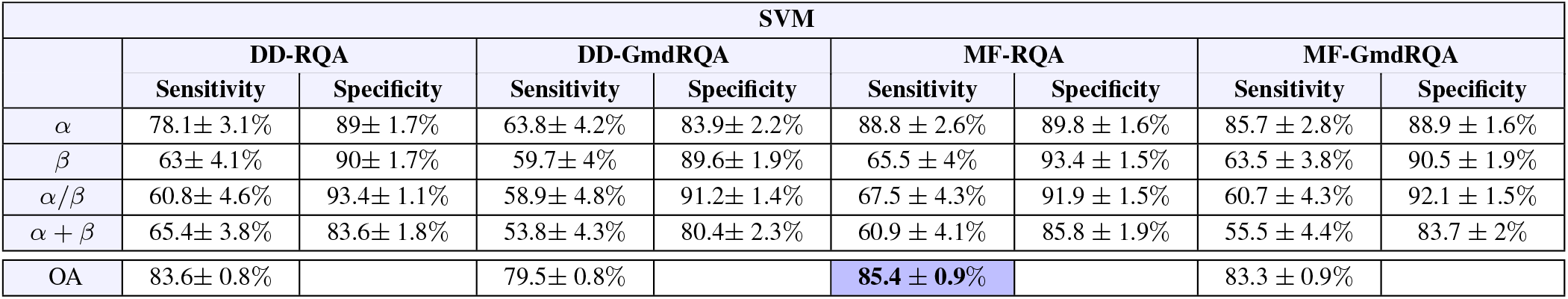
Performance evaluation of (i) the data-driven fine-tuned unidimensional RQA (DD-RQA), (ii) the data-driven fine-tuned generalized multidimensional RQA (DD-GmdRQA), (iii) the most frequent fine-tuned unidimensional RQA (MF-RQA) and (iv)) the most frequent fine-tuned generalized multidimensional RQA (MF-GmdRQA) scheme using Support Vector Machines (SVM). The optimal parameter set for (iii) and (iv) are evaluated in Section 3.1.3.

**Table 2.** Performance evaluation of (i) the data-driven fine-tuned unidimensional RQA (DD-RQA), (ii) the data-driven fine-tuned generalized multidimensional RQA (DD-GmdRQA), (ii) the most frequent fine-tuned unidimensional RQA (MF-RQA) and (iv)) the most frequent fine-tuned generalized multidimensional RQA (MF-GmdRQA) scheme using Fisher’s linear Discriminant Algorithm (FDA). The optimal parameter set for (iii) and (iv) are evaluated in Section 3.1.3.

| FDA |  |  |  |  |  |  |  |  |
| --- | --- | --- | --- | --- | --- | --- | --- | --- |
|  | DD-RQA |  | DD-GmdRQA |  | MF-RQA |  | MF-GmdRQA |  |
|  | Sensitivity | Specificity | Sensitivity | Specificity | Sensitivity | Specificity | Sensitivity | Specificity |
| $\alpha$ | 73.2 ± 4% | 87.1 ± 2.5% | 42.2 ± 4.4% | 93 ± 1.5% | 85.2 ± 2.9% | 92.8 ± 1.4% | 78.3 ± 3.3% | 93.2 ± 1.1% |
| $\beta$ | 53.8 ± 6.8% | 92.6 ± 2.2% | 63.1 ± 3.8% | 85.7 ± 1.8% | 55.3 ± 5.1% | 96.2 ± 1.5% | 64.8 ± 3.8% | 91.3 ± 1.3% |
| $\alpha/\beta$ | 60.9 ± 4.2% | 91.9 ± 1.4% | 55.7 ± 4.7% | 90.9 ± 1.4% | 64.7 ± 4.7% | 91.9 ± 1.5% | 59.6 ± 5% | 92.4 ± 1.4% |
| $\alpha + \beta$ | 68.9 ± 4.1% | 80.2 ± 2.6% | 64.7 ± 4.1% | 72 ± 2.6% | 70.4 ± 3.9% | 77.6 ± 2.7% | 64.3 ± 4.1% | 79 ± 2.4% |
| OA | 82.1 ± 1% |  | 78.2 ± 1% |  | **84.5 ± 0.9%** |  | 83.5 ± 1% |  |

The scheme of RQA with grid search parameter selection has been widely employed in previous studies (Yang *et al.*, 2009, 2010; Olyaee *et al.*, 2016). However, none of the corresponding works report the optimal parameter set-up. Hence, in this work a GS-RQA and a GS-GmdRQA framework are re-implemented. Along these lines, as depicted in Table 3, four different experiments involving the GS-RQA/GmdRQA, DD-RQA/GmdRQA, MF-RQA/GmdRQA and HVG are conducted. In more detail, Experiment 1 indicates that the GS-RQA framework with SVM achieves the highest classification accuracy (86.20%). Furthermore, GS-RQA provides better results than those reported in the respective works of Yang *et al.*, 2009 and Yang *et al.*, 2010. Specifically, the main difference between our proposed GS-RQA and Yang *et al.*, 2009 is the absence of PSI-PRED tool from their architecture. On the other hand, in the work of Yang *et al.*, 2010 PSI-PRED is employed but the number of extracted features is 16, whereas we have extracted 20 features. Moreover, we cannot conclude on the effect of the grid search parameter values on accuracy results since Yang *et al.*, 2010 do not report their grid search parameter values. Nevertheless, the performance of GS-RQA is comparable to MF-RQA with SVM, which reaches an overall classification accuracy of 85.40%. It is clear that GS-RQA improves only slightly the overall classification accuracy (86.20% vs 85.40%) but with the cost of an extremely high running time (427.86 minutes vs 11.15 minutes). Thus, we conclude that MF-RQA is the most efficient architecture in terms of running time complexity with comparable classification accuracy in Experiment 1. Finally, it is also worth mentioning that the best algorithm in terms of computational complexity is MF-RQA. Thereafter, GmdRQA is examined in Experiment 2. As presented, GmdRQA utilizes the minimum number of features that has been so far proposed in the literature (Yang *et al.*, 2009, 2010; Olyaee *et al.*, 2016; Jiang *et al.*, 2019), namely 10, realizing a maximum overall accuracy of 85.30% when GS-GmdRQA with SVM classifier is employed. However, we note that the performance of this scheme comes again with the cost of high computational complexity. Thus, MF-GmdRQA is highlighted as the best trade-off in terms of classification accuracy (83.54%) for the FDA classifier and running time efficiency. The most important outcome of the previous two experiments though is that the overall classification accuracy is similar for the RQA and GmdRQA schemes with the later reducing the computational cost approximately to half, in most cases. Therefore, Experiments 1 and 2 demonstrate that MF-GmdRQA is the most efficient scheme as it yields a high overall accuracy that exceeds the performance of the respective works of Yang *et al.*, 2009 and Yang *et al.*, 2010, utilizing the minimum number of features that has been reported so far considering the RQA framework (Yang *et al.*, 2009, 2010; Olyaee *et al.*, 2016; Jiang *et al.*, 2019). Thus MF-GmdRQA not only reduces time complexity due to its automated parameter estimation, but also improves the memory complexity.

**Table 3.** Summarized predicted quality results for the 25PDB dataset. The minimum computational time as well as the highest achieved accuracy for SVM and FDA per experiment, are indicated with bold letters. Blue highlight refers to the most efficient architecture in terms of classification accuracy and running time complexity per experiment. The classification process is excluded for the measurement of the computational complexity that refers only to the feature extraction procedure.

|  | PSI-PRED | CGR | Method |  |  |  |  |  |  |  | Number<br>of<br>Features | Overall<br>Accuracy |  | Computational<br>Complexity<br>(minutes) |
| --- | --- | --- | --- | --- | --- | --- | --- | --- | --- | --- | --- | --- | --- | --- |
|  |  |  | RQA |  |  | GmdRQA |  |  | HVG | Other |  | SVM | FDA |  |
|  |  |  | GS | DD | MF | GS | DD | MF |  |  |  |  |  |  |
| Yang *et al.*, 2009 |  | X | X |  |  |  |  |  |  |  | 16 | - | 65.08 | - |
| Yang *et al.*, 2010 | X | X | X |  |  |  |  |  |  |  | 16 | - | 81.40 | - |
| Experiment 1 | X | X | X |  |  |  |  |  |  |  | 20 | **86.20** | 84.37 | 427.86 |
|  | X | X |  | X |  |  |  |  |  |  | 20 | 83.60 | 82.14 | **8.18** |
|  | X | X |  |  | X |  |  |  |  |  | 20 | **85.40** | **84.52** | 11.15 |
| Experiment 2 | X | X |  |  |  | X |  |  |  |  | 10 | **85.30** | **84.40** | 232.80 |
|  | X | X |  |  |  |  | X |  |  |  | 10 | 79.51 | 78.23 | **4.38** |
|  | X | X |  |  |  |  |  | X |  |  | 10 | **83.32** | 83.54 | 8.33 |
| Olyaei *et al.*, 2016 | X | X | X |  |  |  |  |  |  | X | 24 | - | **90.08** | - |
| Jiang *et al.*, 2019 | X | X | X |  |  |  |  |  | X | X | 30 | **95.33** | - | - |
| Experiment 3 | X | X | X |  |  |  |  |  | X |  | 37 | 86.16 | 85.92 | 430.76 |
|  | X | X |  | X |  |  |  |  | X |  | 37 | **85.95** | **86.69** | **11.08** |
|  | X | X |  |  | X |  |  |  | X |  | 37 | **86.20** | 86.25 | 14.06 |
| Experiment 4 | X | X |  |  |  | X |  |  | X |  | 27 | 85.37 | 87.08 | 235.71 |
|  | X | X |  |  |  |  | X |  | X |  | 27 | **84.39** | **86.30** | 7.28 |
|  | X | X |  |  |  |  |  | X | X |  | 27 | **86.27** | **87.23** | 11.24 |

Considering the works of Olyaee *et al.*, 2016 and Jiang *et al.*, 2019, RQA is combined with other feature extraction and time series analysis techniques that are not examined in the herein work. Particularly, Olyaee *et al.*, 2016 combine RQA and Complex Networks whereas Xu *et al.*, 2017 combined the multiscale coarse-grained RQA with HVG. In our work, HVG is also employed and combined with all the aforementioned RQA and GmdRQA schemes in Experiment 3 and 4. The main benefit of HVG against RQA is the absence of hyper-parameter tuning that results in low running time complexity. In particular, our implementation requires 2.9 minutes to reproduce the HVG framework proposed by Zhao *et al.*, 2018. As depicted in Experiment 3, the combination of HVG with GS-RQA, DD-RQA and MF-RQA slightly improves the overall classification accuracy with the HVG and DD-RQA being the optimal scheme in terms of classification accuracy (86.69%) and time and memory complexity. The performance of the combination of HVG and GmdRQA frameworks is then presented in Experiment 4. The combination of HVG and DD-GmdRQA achieves similar overall classification accuracy with the rest architectures, however, it is more favorable in terms of computational complexity.

Compared to the best performing scheme of this work, i.e. DD-GmdRQA and HVG, the work of Olyaee *et al.*, 2016 achieves higher overall accuracy, namely 90.08%, utilizing 24 features that derive from the combination of RQA and Complex Networks. However, the tuning of the RQA hyperparameters is performed in a grid search manner which is extremely time consuming based on the experiments presented in this work for GS-RQA. The same assertion can be also made for the work of Jiang *et al.*, 2019 although the stage of parameter selection and definition is not stated in their work. Specifically, they utilize multiscale coarse-grained RQA (Xu *et al.*, 2017) along with HVG and achieve an overall accuracy of 95.33%. However, multiscale coarse-grained RQA considers the spatial proximity of the phase space of time series adding an extra hyperparameter to the system and hence further escalates the time complexity. In addition, the work of Jiang *et al.*, 2019 implements an extremely time consuming classification procedure. In particular, they perform a leave-one-out cross-validation using an SVM classifier with a grid search parameter selection of the regularization parameter *C* and kernel width parameter *γ* that can take all positive values log-scaled in the range [2^−10^, 2^10^]. On the contrary, the implementation of FDA in our work is performed in an automated fashion and requires merely 2.98 seconds, whereas our proposed SVM classification scheme requires 24.87 seconds for the exact same setup described in Section 3.2.

## 4 Conclusion

In this work, we designed and implemented novel data-driven protein structure prediction architectures based on the representation of secondary structure data in higher-dimensional phase spaces using RQA, GmdRQA, and a combination of HVG with the two aforementioned techniques. In particular, the herein proposed work addresses the problem of efficient time delay embedding parameter selection which has a significant positive impact on the overall classification accuracy and the running time complexity. Four efficient data-driven evaluation approaches are suggested, namely DD-RQA, DD-GmdRQA, MF-RQA, and MF-GmdRQA. The experimental evaluation on real data revealed the superiority of the HVG & DD-GmdRQA-based framework in extracting and exploiting the underlying temporal dynamics of the data generating processes, resulting in lower time and memory complexity in terms of feature multitude, when compared against the state-of-the-art. An extension of this work will consider a GmdRQA framework that processes directly on the primary amino acid protein sequence without employing intermediate tools such as PSI-PRED. Lastly, a future goal is to predict the three-dimensional structure of the proteins with deep learning algorithms, exploiting the graphs exported with CGR.

